# Luminal MCF-12A and myoepithelial-like Hs 578Bst cells form bilayered acini similar to human breast

**DOI:** 10.1101/248765

**Authors:** Anne Weber-Ouellette, Mélanie Busby, Isabelle Plante

## Abstract

The mammary gland is a complex organ, structured in a ramified epithelium supported by the stroma. The epithelium’s functional unit is the bilayered acinus, made of a layer of luminal cells surrounded by a layer of basal cells mainly composed of myoepithelial cells. The aim of this study was to develop a reproducible and manipulable three-dimensional co-culture model of the bilayered acinus *in vitro* to study the interactions between the two layers. Two different combinations of cell lines were co-cultured in Matrigel: SCp2 and SCg6 mice cells, or MCF-12A and Hs 578Bst human cell lines. Cell ratios and Matrigel concentration were optimized. The resulting acini were analysed by confocal microscopy using epithelial (E-cadherin) and myoepithelial (α-smooth muscle actin) markers. SCp2 and SCg6 cells formed distinct three-dimensional structures, whereas MCF-12A and Hs 578Bst cells formed some bilayered acini. This *in vitro* bilayered acini model will allow us to understand the role of interactions between luminal and myoepithelial cells in the normal breast development.

## 2.3.2 Introduction

The human mammary gland consists of two compartments: the stroma and the epithelium. The mammary epithelium is organized in a ramified lobulo-alveolar system. In humans, 15 to 20 lobes in each breast converge in ducts toward the nipple. Each lobe is itself sub-divided in lobules, and each lobule consists of many acini grouped together (Hassiotou & Geddes, 2013). The acinus is considered to be the functional unit of the mammary gland. The whole ramified lobulo-alveolar system of the epithelium consists of a central lumen bordered by an inner layer of luminal cells surrounded by an outer layer, mainly comprised of myoepithelial cells. The epithelium is separated from the mammary stroma by a basement membrane (Su, Shankar et al., 2011). In the acini, the myoepithelial cells form a basket-like network surrounding the luminal cells (Sopel, 2010).

The stroma surrounding the breast epithelium is comprised of extracellular matrix (ECM), of mesenchymal and immune cells, as well as of blood and lymphatic vessel cells (Weigelt, Ghajar et al., 2014). The basement membrane is a type of specialized ECM mainly composed of type IV collagen and laminin-1 (Leblond & Inoue, 1989). In addition to physically supporting the epithelium, the stroma’s components transmit signalling cues to the epithelium including through transmembrane receptors, such as integrins, to the cytoskeleton of the cell, which ultimately impinges on chromatin and nuclear function to maintain tissue integrity (Bissell, Hall et al., 1982). Hence, the environment in which cells grow, or in which cells are cultured *in vitro*, impacts their morphological organization and tissue-specific functions (Weigelt et al., 2014). Traditionally, cultured cells are grown in a two dimensional (2D) monolayer. They adhere and grow on a flat surface, and adopt a flatter and more stretched out phenotype compared to *in vivo*. This abnormal cell morphology alters cellular processes such as cell proliferation, differentiation, apoptosis, as well as gene and protein expression (Tibbitt & Anseth, 2009). Consequently, cells cultured in 2D may not behave as they would in tissues (Huh, Hamilton et al., 2011).

To overcome the limitations of traditional 2D cell culture and to adequately mimic the *in vivo* microenvironment, cells must be cultured in three dimensions (3D). This type of culture allows cells to freely assemble in multidimensional structures, called spheroids, using a scaffold/matrix or in a scaffold-free manner (Edmondson, Broglie et al., 2014). By adopting this *in* vivo-like 3D relationship to each other, cells better can form cell-cell and cell-ECM interactions, establish appropriate cell-signalling pathways to maintain tissue function and mimic the cellular processes occurring in the human body (Edmondson et al., 2014). This difference between 2D and 3D culture regarding cellular processes and response, through the modulation of gene expression, has been observed several times (Chitcholtan, Asselin et al., 2013; Mabry, Payne et al., 2016; Pineda, Nerem et al., 2013). In fact, distinct patterns in gene expression profiles between tissue samples and cell lines of varying phenotypes demonstrated adaptation of cells to their culture microenvironment (Birgersdotter, Sandberg et al., 2005; Kenny, Lee et al., 2007).

Most 3D cell culture systems mimicking mammary gland acini are monocultures, using only the luminal cells (Debnath, Muthuswamy et al., 2003; Lee, Kenny et al., 2007; Marchese & Silva, 2012). Yet, to thoroughly recapitulate the histological complexity of the normal human mammary gland, luminal cells must crosstalk with EMC or scaffolds, but also with myoepithelial cells (Radisky, Hagios et al., 2001). *In vivo*, myoepithelial cells interact physically and by paracrine signalling with the layer of luminal epithelial cells, and are critical for the proper polarisation of the luminal cells (Gudjonsson, Ronnov-Jessen et al., 2002). Heterotypic models of the human mammary gland have been developed using luminal MCF-10A cells, adipocytes and human fibroblasts in a mixture of laminin-rich basement membrane extract (lrBME) matrix/collagen on porous silk protein scaffold (Wang, Sun et al., 2010). While this model represents progress towards an *in vitro* acinus-like structure composed of multiple cell types, it uses complex matrices and scaffolds, whereby myoepithelial cells are absent. Bilayered acini composed of a mix of purified human luminal and myoepithelial cells isolated from normal mammary glands have also been developed (Gudjonsson et al., 2002). However, because they are formed using primary cultures, they are hardly genetically manipulable. There is thus still a great need for a simplified, optimized, genetically manipulable, reproducible and physiologically relevant model to recapitulate the normal structure of the functional unit of the human mammary gland – the bilayered acinus (Campbell & Watson, 2009).

This study aimed to develop a model representing the breast bilayered acini that can be genetically manipulated and easily reproduced by using cell lines. Here, two combinations of cell lines were tested: 1) the human non-tumorigenic luminal and myoepithelial-like cells MCF-12A and Hs 578Bst; and 2) the murine non-tumorigenic luminal and myoepithelial cells SCp2 and SCg6 (Desprez, Roskelley et al., 1993).

## 2.3.3 Results

### 2.3.3.1 MCF-12A and Hs 578Bst cells showed more differentiated phenotypes than SCp2 and SCg6

We first confirmed that the selected cell lines were representative of luminal and myoepithelial cells by evaluating the expression of various markers (Supplementary figure 1). While MCF-12A and Hs 578Bst cells expressed markers typical for luminal and myoepithelial cell types respectively, some markers were present in both SCp2 and SCg6 cells (Supplementary figure 1).

### 2.3.3.2 MCF-12A and SCp2 cells embedded in Matrigel form spheroid structures

We then wanted to confirm that human mammary gland luminal MCF-12A and SCp2 cell can form acini-like structures, as reported in the literature (Marchese & Silva, 2012; Talhouk, Mroue et al., 2008). To do so, MCF-12A were embedded in a 3D matrix consisting of cell culture medium and basement membrane extract, commercially known as Matrigel. After 24 hours in culture, cells already formed small clusters of cells (Figure 1A). After 4 days, small and rounded spheroid structures could be observed (Figure 1B). After 10 days, the spheroids were still present, but were much bigger in size, and the cells that constituted each spheroid could be distinguished (Figure 1C). After 14 days in culture, spheroids maintained their size and more defined round structure started to form in some of the spheroids, similar to the acini of the human mammary gland (Figure 1D). Likewise, SCp2 cells could form acini-like structures with a lumen (Supplementary figure 2A and B).

**Figure 1:**
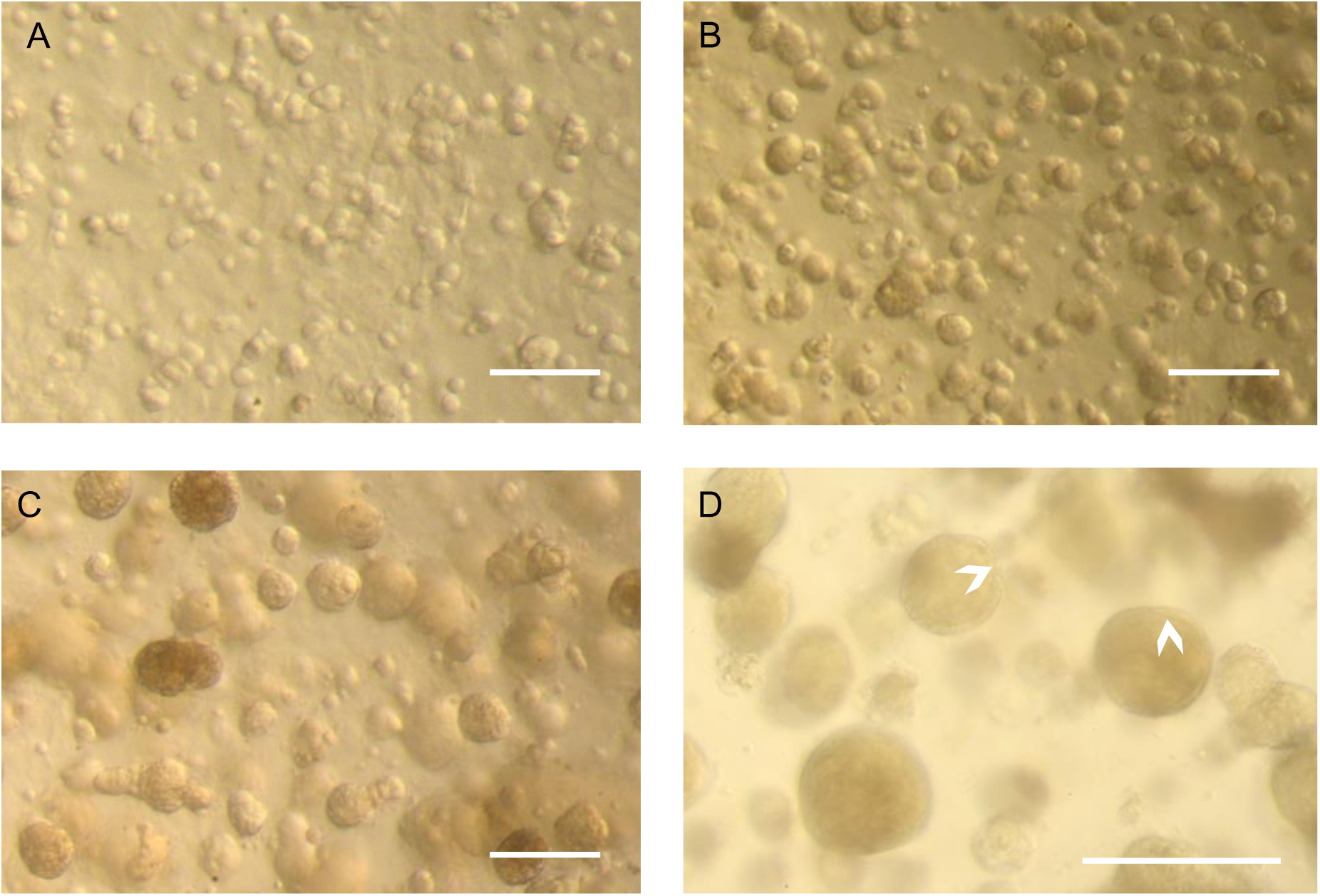
MCF-12A cell embedded in Matrigel gradually form acini-like structures. (**A-D**) Optical microscopy images of MCF-12A. Cells appeared in small clusters after 24h (**A**). They gradually formed acini-like spheroids after 4 days (**B**), which grow in size after 10 days (**C**). 14 days after being embedded in Matrigel, spheroids preserved their size and some of them were more defines, with elea) ed ges (**D, arrowheads**). Scale bars: 250 μm.

### 2.3.3.3 MCF-12A cells maintained in Hybri-Care culture medium conserved an acini-like structure

Because Hs 578Bst and MCF-12A cells are typically grown in different media, and because Hs 578Bst cells are more difficult to maintain than MCF-12A cells, we wanted to insure that MCF-12A could still form acini-like structures in Hs 578Bst cells culture media. We thus compared spheroids formed by MCF-12A cells embedded in Matrigel and maintained with either Hybri-Care medium (ATCC^®^ 46-X^™^) or phenol red-free DMEM/F12 medium, the media typically used for Hs 578Bst and MCF-12A cells, respectively. While some MCF-12A spheroids showed a defined spherical structure when maintained in DMEM/F12 medium, smaller less defined structures, characterized by a certain looseness in the structure, were also present (Figures 2A-C). These spheroids also displayed what looks like cellular degradation at their edges (Figure 2A-C). Interestingly, when Hybri-Care medium was used to dilute the Matrigel and to maintain embedded cultures, the cells formed bigger, rounder and more defined acini-like spheroids. These structures were more compact and there was no visual evidence of cellular degradation (Figure 2D-F). These observations suggested that MCF-12A cells can form acini-like structures even more efficiently in Hybri-Care medium. Because SCp2 and SCg6 cells are cultured in the same medium, these optimizations were not necessary.

**Figure 2:**
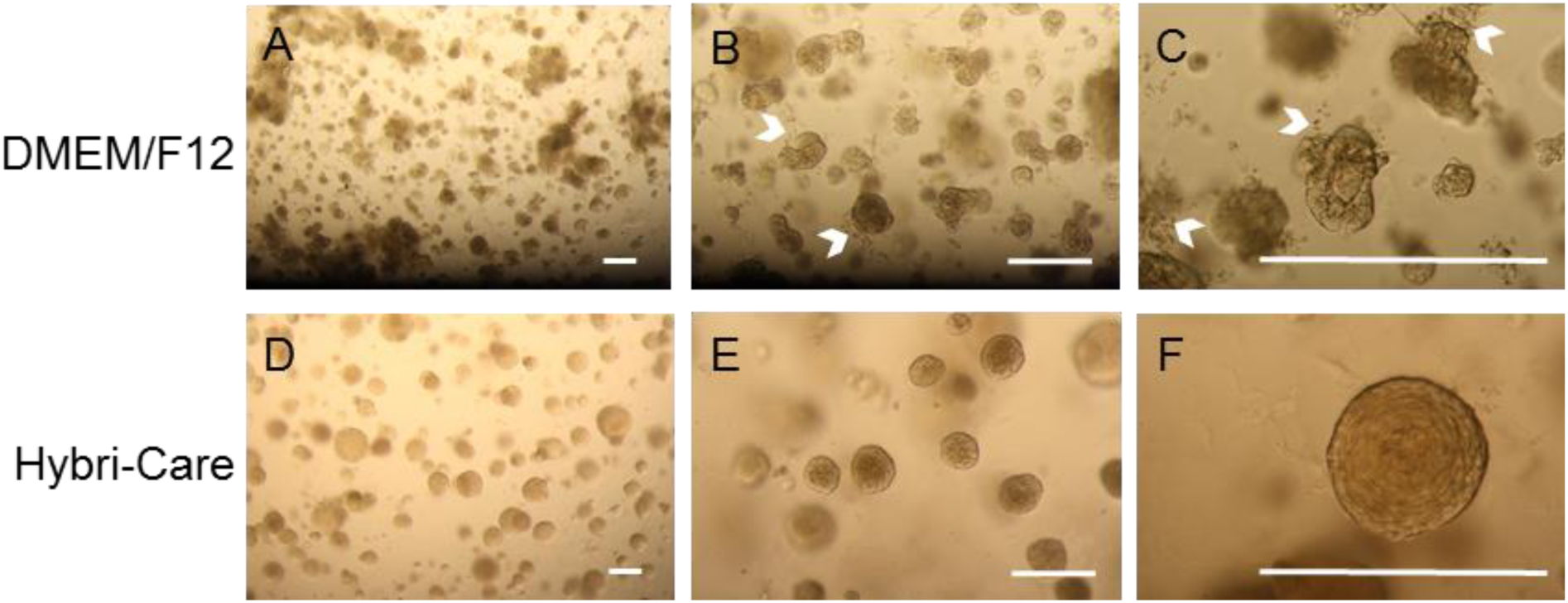
Hybri-Care culture medium promotes the formation of more defined acini-like structure than DMEM/F12 culture medium. (**A-F**). Optical microscopy images of MCF-12A cell embedded in Matrigel after 14 days of culture. When using DMEM/F12 medium to maintain cultures, resulting multicellular structures seem smaller, less defined and looser (**A-C, arrowheads**) than cells maintained in Hybri-Care medium (**D-F**). In hybrid-Care medium, acini are bigger, rounder, more defined and more compact. Scale bars: 250 μm.

To ensure that these spheroids maintained the lumen characteristic of acini, they were analyzed by confocal microscopy. The acini presented lower cell density at their center, suggesting the gradual apoptotic clearance of cells creating a lumen-like cavity (Figure 3). These results were confirmed by cryosections of MCF-12A acini, either stained using Masson's trichrome staining (Figure 3C) or immunolabeled with β-catenin-specific antibody (Figure 3D). Adherens junctions were formed between luminal cells as demonstrated by the expression of both E-cadherin and β-catenin (Figure 3A, B and D)

**Figure 3:**
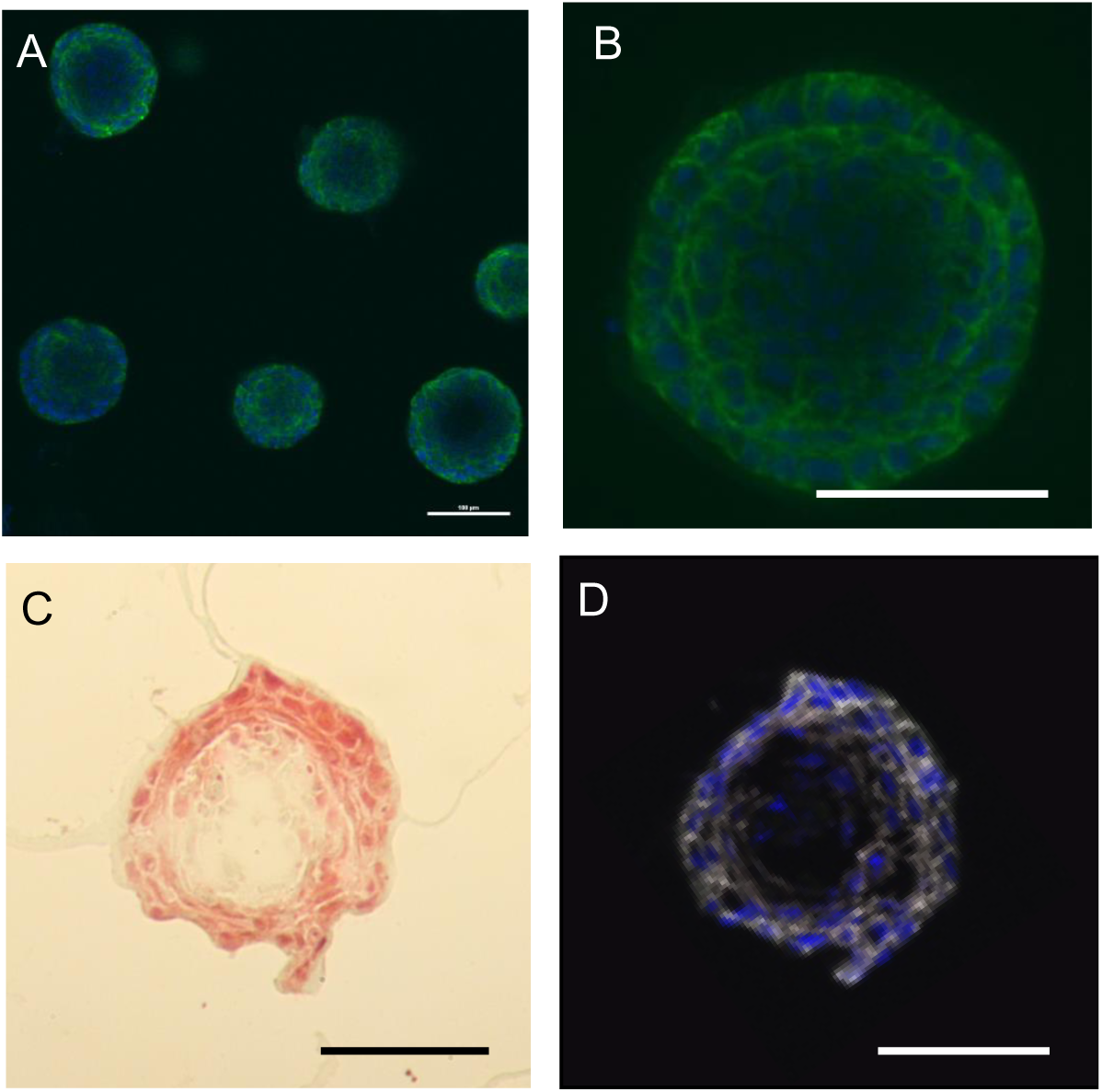
Presence of a lumen-like cavity in the center of cryosectionned MCF-12A acini. (**A, B**). Confocal microscopy images of acini immunolabeled with an E-cadherin (adherens junctions) specific antibody (green). Nuclei are stained with DAPI (Blue). (**C**) Optical microscopy image of an acini cryosection stained with Masson’s trichrome staining. (**D**) Confocal microscopy image of an acini cryosection immunolabelled with a β-catenin (adherens junctions) specific antibody (white). Nuclei are stained with DAPI. Scale bars: 100 μm.

### 2.3.3.4 Dilution of Matrigel to different concentrations impacts spheroid structure and immunolabeling

To optimize the immunolabeling of the acini-like spheroids, without compromising their 3D structure, we then assessed which Matrigel concentration is most favorable to support round acini formation and to facilitate sufficient antibody penetration in the matrix for decent immunofluorescence imaging. MCF-12A cells were embedded in Matrigel diluted with Hybri-Care medium to achieve concentrations of 50%, 75% and 100% of Matrigel (Figure 4). In wells containing 50% Matrigel, the matrix almost completely liquefied during the immunolabeling procedure, resulting in the loss of many acini during staining procedure. The remaining acini fell to the bottom of the microwell, forming flat structures difficult to properly label with antibodies (Figure 4A and B). On the opposite, in wells containing 100% of Matrigel, the matrix remained rigid in its consistency, resulting in unspecific labeling and smaller acini (Figure 4E and F). Finally, in wells containing a concentration of 75% of Matrigel, the matrix had a soft consistency, but preserved its rigidity through washes, and immunolabeling was specific (Figure 4C and D).

**Figure 4:**
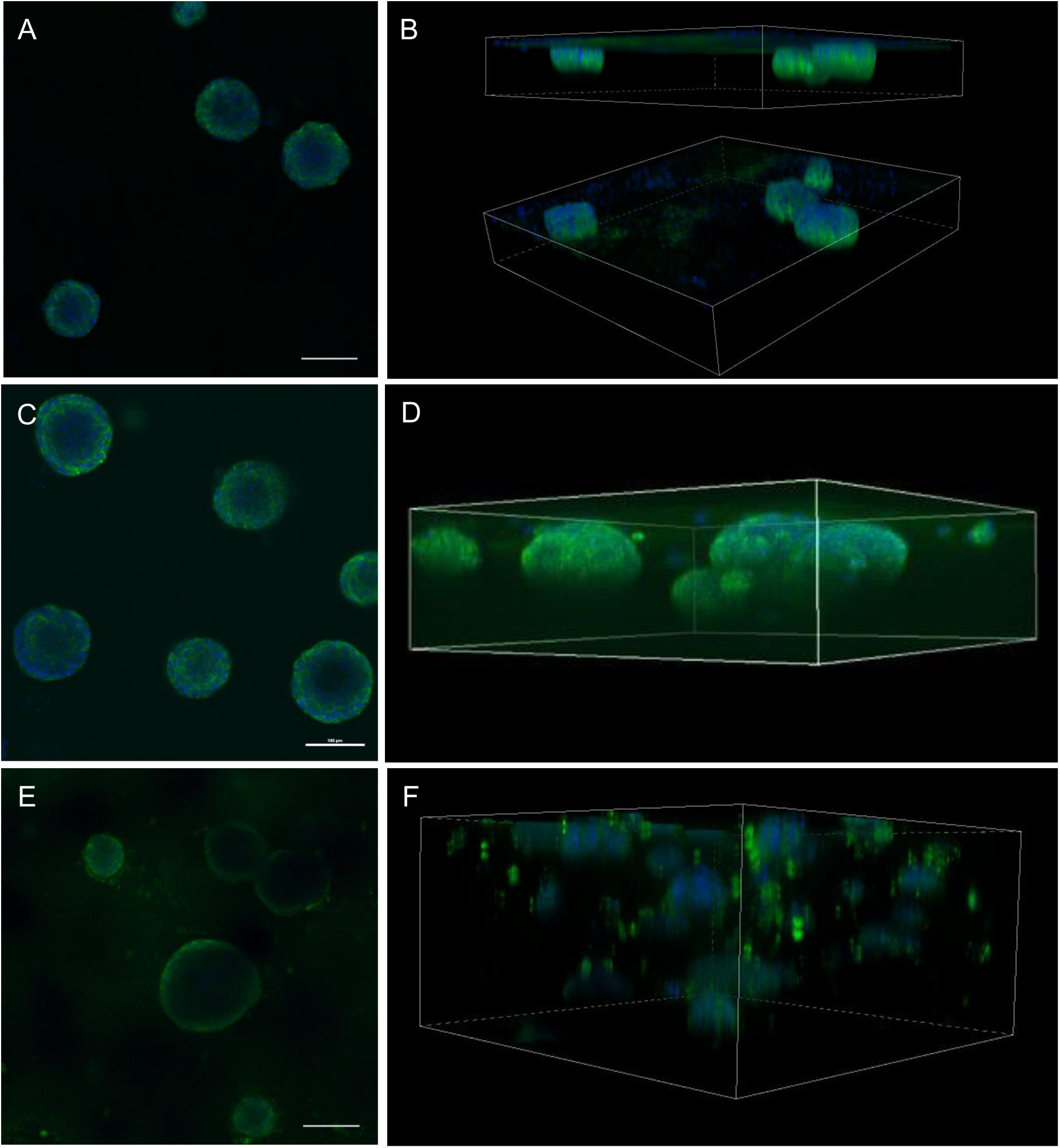
Matrigel at a concentration of 75°% is optimal for 3D culture of MCF-12A cells. (**A, B**). Representative single confocal images (**A, C, E**) or serial Z-stacks (**B, D, F**) of MCF-12A cells embedded in 50% (**A, B**), 75% (**C, D**) or 100% (**E, F**) Matrigel and immunolabeled with an E-cadherin antibody (green). Nuclei are stained with DAPI (blue). Scale bars: 100 μm.

### 2.3.3.5 MCF-12A and Hs 578Bst self-organized in spheroids resembling bilayered-like acini

Once culture conditions were optimized for MCF-12A, we co-cultured luminal MCF-12A cells with myoepithelial-like Hs 578Bst cells, as well as SCp2 and SCg6 cells. Similar to when MCF-12A cells were in 3D monoculture (Figure 5A and B), acini-like structures formed when MCF-12A and Hs 578Bst were simultaneously embedded in Matrigel (Figure 5C and D). In the co-culture, however, a second layer of cells seemed to surround some of the acini, suggesting that bilayered acini were formed with both types of cells (Figure 5D, arrowhead). Hs 578Bst cells, when cultured alone, were not able to form spheroids (Figure 5E). On the opposite, SCg6 cells could form spheroids alone or in co-culture with SCp2 (Supplementary figure 2C-E).

**Figure 5:**
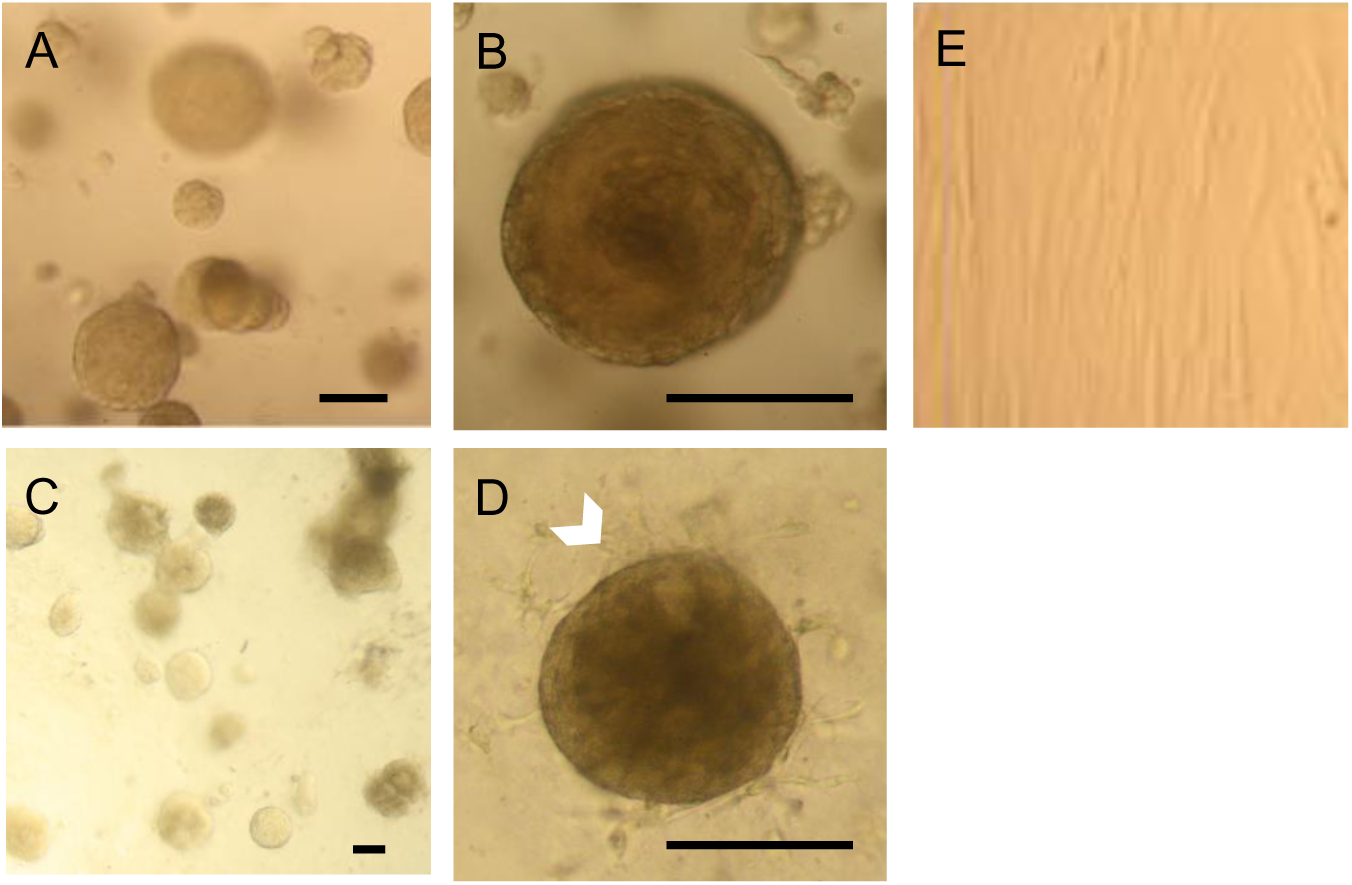
MCF-12A cells co-cultured with Hs 578Bst cells produce acini-like structures when embedded in Matrigel. (**A, B**) Representative optical microscopy images of MCF-12A cells embedded in Matrigel. (**C, D**) Optical microscopy images of MCF-12A cells co-cultured with Hs 578Bst cells showing acini-like structures. A second layer of cells seems to be surrounding the first layer in some of the acini (arrowhead), suggesting that myoepithelial cells formed bilayered acini with the luminal cells. (**E**) Optical microscopy image of Hs 578Bst cells embedded in Matrigel. Scale bars: 100 μm.

To confirm that these acini were indeed bilayered, we performed immunolabeling using distinct epithelial (E-cadherin) and myoepithelial (smooth muscle actin, SMA) markers. While some spheroids with lumen were present when SCp2 and SCg6 cells were co-cultured in 3D, most structures were undefined spheroids (Supplementary figure 2F). Importantly, none of those structures were bilayered. However, when luminal MCF-12A cells were co-cultured with myoepithelial-like Hs 578Bst cells, bilayered acini were produced (Figure 6). These acini were composed of an inner layer of luminal cells surrounded by a discontinuous basket-like network of myoepithelial-like cells (Figure 6B). Moreover, the truncated view allowed us to distinguish a lumen in the center of the acini (Figure 6C). From these results, we can conclude that it is possible to create a 3D model of a human bilayered acini *in vitro*, using cell lines.

**Figure 6.**
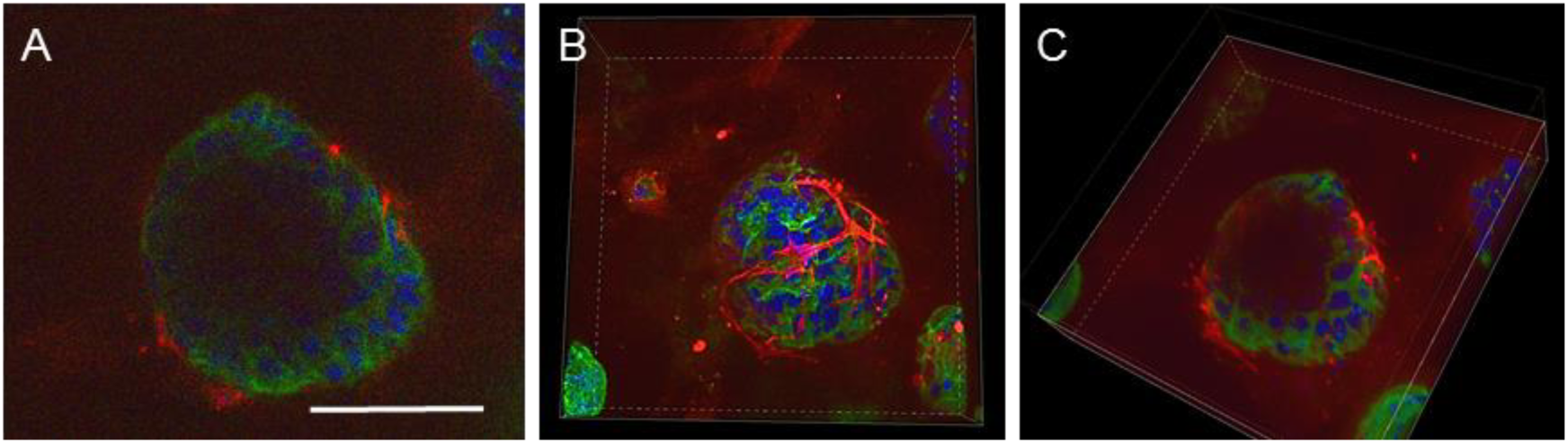
MCF-12A and Hs 578Bst co-cultured cells form bilayered acini in Matrigel. (**A-C**) Confocal microscopy images of MCF-12A and Hs 578Bst cells forming a 3D bilayered acini when co-cultured in Matrigel. Acini were immunolabeled with an E-cadherin specific antibody (green) and a SMA specific antibody (red). Nuclei are stained with DAPI (blue). Representative single confocal image (**A**), serial Z-stack (**B**) and a truncated view of a Z-strac (**C**). The myoepithelial-like cells Hs 578Bst have a stellate phenotype, forming a discontinuous basket-like network around the MCF-12A luminal cells. The truncated view suggest the presence of a lumen in the center of the bilayered acini. Scale bar: 100 μm.

## 2.3.4 Discussion

To fully understand the mechanisms that lie behind breast cancer, we first need to understand how a healthy mammary gland functions. Developing a surrogate model of the normal human breast that mimics the architecture and function of the actual organ will help increase our understanding of how breast tissue develops and how specific deregulations, of intercellular junctional complexes for example, influence carcinogenesis. Many techniques have been explored to model and study the human mammary gland *in vitro*. Malignant as well as non-malignant mammary cells have traditionally been studied as monolayer on plastic cell culture dishes, thereby losing their morphological organization and tissue-specific function (Weigelt et al., 2014). Fortunately, progress in tissue engineering and biomaterials have provided researchers with innovative techniques that are now allowing to explore the possibilities of 3D culture, thus bridging the gap between *in vitro* monolayer cell culture models and expensive *in vivo* whole-animal systems (Campbell & Watson, 2009). 3D culture systems promote expression of tissue-specific functions and cellular processes by allowing cells to self-assemble and to receive cues from their neighbouring cells and the surrounding extracellular matrix, which cannot be achieved when cells are plated on plastic cell culture dishes in 2D (Campbell & Watson, 2009; Weigelt & Bissell, 2008). 3D models are particularly useful for the study of protein and gene functions, along with signaling pathways in a physiologically relevant context (Weigelt & Bissell, 2008).

Here we report the production of a relevant 3D heterotypic co-culture model of the functional unit of the mammary gland, the bilayered acinus, consisting of two different cell lines, human myoepithelial-like Hs 578Bst and luminal MCF-12A. These spherical bilayered structures consisted of a lumen surrounded by an inner layer of luminal cells enveloped by a basket-like network of myoepithelial cells, similar to what is typically observed *in vivo*.

### 2.3.4.1 The importance role of myoepithelial cells

There has been a wide range of 3D culture models, using Matrigel based matrices, developed in an attempt to replicate the epithelium of the human mammary gland *in vitro*. For instance models using non-tumorigenic human mammary luminal cells lines, such as MCF-10A (Debnath et al., 2003), HMT 3522 S1 (Anders, Hansen et al., 2003; Lee et al., 2007) or MCF-12A (Marchese & Silva, 2012) have been reported. When grown in Matrigel, all these cells lines were able to form organized spheroids with a central lumen, similar to breast acini morphology. In opposite, tumorigenic human mammary luminal cells lines form disorganized, proliferative and nonpolar colonies (Kenny et al., 2007; Lee et al., 2007). While these models brought important insights on the structure and the polarization of acini, and the lack of defined structures for breast cancer cells, these models fail to consider the crucial role of myoepithelial cells in the formation of a polarized epithelium *in vivo*.

For many years, myoepithelial cells were mostly ignored in mammary gland studies, as it was thought that their role was limited to transportation and ejection of milk during lactation. We now know that myoepithelial cells are crucial for the proper differentiation and function of the epithelium. Among their functions, they allow paracrine regulation and cross-talk within the epithelium while playing an active role in tissue remodeling and polarization of luminal cells. Such functions are critically dependent on bi-directional communication between myoepithelial and luminal cells (Dickson & Warburton, 1992; Rudolph-Owen & Matrisian, 1998; Talhouk et al., 2008; Warburton, Mitchell et al., 1982). Moreover, a growing body of evidence demonstrates that the tumor microenvironment plays a critical role in cancer progression (Ghajar & Bissell, 2008). Whether cancer cells induce remodeling of the architecture and/or changes in tissue architecture promote cell tumorigenicity is unclear. It is likely that gene expression is at least in part dictated by the interactions between a cell and the stromal elements, including stromal cells, proteins of the ECM and other soluble factors (Ghajar & Bissell, 2008). As a result, ECM is considered as an active participant, rather than a passive bystander, in cellular differentiation. Conversely, cells contribute to the formation of the epithelial microenvironment by producing components of the basement membrane such as collagen, laminins and fibronectin (Warburton et al., 1982). Because they lie on the epithelial side of the basement membrane, myoepithelial cells are uniquely positioned to accomplish most of the interactions between the epithelium, basement membrane and the ECM. Myoepithelial cells are crucial mediators of ductal elongation and invasion within the stroma, as they secrete proteins and molecules required for the remodeling of the ECM such as maspin, amyloid beta-protein precursor/ protease nexin-II (APP/PNII) and matrix metalloproteinases (MMPs) (Dickson & Warburton, 1992; Rudolph-Owen & Matrisian, 1998; Warburton et al., 1982). As a result, dysregulation of myoepithelial cells functions has been associated with the loss of a polarized epithelium, developmental defects and tumorigenesis (Plante & Laird, 2008; Radisky & Radisky, 2010; Runswick, O'Hare et al., 2001; Streuli, Schmidhauser et al., 1995). Accordingly, myoepithelial cells are often considered natural tumor suppressors due to their ability to build a physical and chemical barrier against uncontrolled growth, tumor cell invasion and angiogenesis (Barsky & Karlin, 2005; Sternlicht, Kedeshian et al., 1997). Therefore, to fully understand both the normal development of mammary gland and breast tumor progression, as well as to study more specifically the direct relationship between luminal and myoepithelial cells of the acinus (Gudjonsson, Adriance et al., 2005), heterotypic models in which human mammary myoepithelial cells are introduced in a human mammary luminal cell culture must be developed.

### 2.3.4.2 Luminal and myoepithelial-like commercial cell lines can form bilayered acini, similar to complex 3D structures from primary cultures

A few studies have been published with 3D models composed of more than one cells types. For instance, human luminal MCF-10A cells were co-cultured with primary culture of human mammary fibroblasts (Krause, Maffini et al., 2008). In another study, primary cultures of human luminal and myoepithelial cells were co-cultured (Carter, Gopsill et al., 2017). In an even more complex model, human luminal MCF-10A cells were co-cultured with primary cultures of human mammary fibroblasts and adipose-derived stem cells (Wang et al., 2010). Cells in these co-culture models displayed more differentiated morphological phenotypes and functional activity than in less complex monocultures (Wang et al., 2010). Likewise, these co-cultures facilitated the study of cellular crosstalk in the breast (Carter et al., 2017). Notably, most of these studies used primary cultures of breast cells. It is believed that primary cells in a 3D mammary gland model enable more physiologically relevant studies such as lineage commitment and plasticity (Campbell, Davidenko et al., 2011), and that normal *in vivo* signaling pathways is more preserved compared to immortalized cell lines (Ip & Darcy, 1996). However, some of the downsides of these models are the increased complexity of working with primary cells, the heterogeneity of the samples and the difficulties to genetically manipulate the cells. On the opposite, commercially available cell lines represent a more homogenous population that can easily be genetically modified to isolate the role of particular proteins or pathways (Kaur & Dufour, 2012; Wang et al., 2010). As such, our model using cell lines represent a more reproducible and manipulable model to study the role of the bi-directional cross-talk between luminal and myoepithelial cells within the mammary epithelium. Moreover, because MCF-12A and Hs 578Bst cell lines are commercially available, researchers around the world will be able to use and even improve this model.

### 2.3.4.3 Limitations of our model

Although we successfully obtained bilayered acini, about 50% of the acini formed when we co-cultured luminal and myoepithelial cell were bilayered, while the other 50% was formed of luminal cells only. A possible explanation lies in the use of Matrigel. Indeed, it has been reported that co-culturing isolated myoepithelial and luminal cells in type I collagen based matrix promoted their re-arrangement in bilayered acini, while Matrigel did not (Carter et al., 2017; Gudjonsson et al., 2002). Matrigel is a laminin-rich basement membrane extract. It has been demonstrated that laminin is required for adequate luminal cell polarization in the mammary gland (Gudjonsson et al., 2002), and that myoepithelial cells produce an important amount of this laminin *in vivo* (Deugnier, Moiseyeva et al., 1995). Consequently, in a Matrigel laminin-rich matrix, as laminin is already present, there is no incentive for luminal cells to coalesce with myoepithelial cells in bilayered structures. On the opposite, in a type I collagen based matrix in which no laminin is present, the luminal cells require the laminin produced by myoepithelial cells to promote their assembly into co-units. Yet, we still managed to produce bilayered acini of luminal and myoepithelial cell lines embedded in a Matrigel matrix in our model. Other ECM, such as artificial scaffold or type I collagen based matrix, might help increasing the ratio of bilayered versus luminal cells-only acini. Increasing the myoepithelial to luminal cell ratio in co-culture might also favour the formation of bilayered acini.

The type of cell lines used also seem to a have limitations. In the experiment herein, murine luminal cells SCp2 and myoepithelial cells SCg6 did not interact to form bilayered acini when co-cultured in Matrigel. Both cell lines either formed monolayer acini or irregular structures, but did not coalesce (Supplementary figure 2). This might be explained by the fact that SCp2 and SCg6 cells have less differentiated phenotypes. In fact, Western blot analysis showed that both SCp2 and SCg6 expressed luminal marker E-cadherin and myoepithelial marker cytokeratin 14 (Supplementary figure 1). Moreover, they showed a high plasticity when cultured in 2D, suggesting stem-like properties (data not shown). Finally, while MCF-12A and Hs 578Bst cells have more differentiated phenotypes and are considered as non-tumorigenic, they remained transformed cells. Therefore, signaling and protein expression is surely different than those of *in vivo* luminal and myoepithelial cells. Nevertheless, the fact that they form bilayered acini *in vitro* represents a first step toward more complex, manipulable, reproductive and representative *in vitro* models to mimic the bilayered mammary gland epithelium and study the role of bidirectional communication between the two layers of epithelial cells.

## 2.3.5 Conclusion

To our knowledge, this study is the first to utilize two types of human mammary cell lines cultured together in Matrigel to form bilayered acini, offering significant advantages over previously described models that used monocultures of cell lines or co-culture of primary cells. The critical advantage of our model remains the use of commercially available cell lines that ensure a manipulable, reproducible and physiologically relevant human mammary gland model. Our model will allow the study of the critical role of myoepithelial cells, and of the interactions between myoepithelial and luminal cells in mammary gland development and during breast cancer progression.

## 2.3.6 Materials and methods

### 2.3.6.1 Cell lines

MCF-12A cells (ATCC^®^ CRL-10782) and Hs 578Bst cells (ATCC^®^HTB-125) were purchased at ATCC (ATCC, Manassas, VA). SCp2 and SCg6 cells were a gift from Calvin Roskelley (UBC). MCF-12A cells were maintained in phenol red-free Dulbelcco’s modified Eagle’s medium Ham’s F12 (DMEM/F12) culture medium (21041025, ThermoFisher Scientific, Rockford, Illinois, USA) supplemented with 5% (v/v) horse serum (ThermoFisher Scientific, 16050-122), human Epidermal Growth Factor (hEGF) recombinant (20 ng/ml) (PHG0311, Invitrogen, Waltham, MA, USA), hydrocortisone (500 ng/ml) (H0888, Sigma-Aldrich, Oakville, Ontario, Canada), insulin (10 μg/ml) (Sigma-Aldrich, C8052), cholera toxin (100 ng/ml) (Invitrogen, 12585014) and propagated according to ATCC guidelines. Hs 578Bst cells were maintained in Hybri-Care medium (ATCC 46-X^™^) supplemented with 10 % (v/v) activated fetal bovine serum (FBS) (098150, Wisent Bioproducts, Saint-Jean-Baptiste, Quebec, Canada) and mouse Epidermal Growth Factor (Epidermal Growth Factor from murine submaxillary gland, 30ng/mL) (Sigma-Aldrich, E4127) and propagated according according to ATCC guidelines. SCp2 and SCg6 cells were grown in DMEM/F12 medium (Sigma-Aldrich, D2906) supplemented with insulin (5 μg/ml) and FBS (5% v/v).

### 2.3.6.2 Western blot analysis

Cell monolayers were washed twice with PBS before the addition of lysis buffer (Tris 50 mM, NaCl 150 mM, 0.02% sodium azide, 0.1% SDS, 1% Nonidet P40, 0.5% sodium deoxycholate, pH: 8) supplemented with NaF 1.25 M, NaVO3 1 M and Halt Protease and phosphatase cocktail inhibitor (Fisher Scientific Canada). Cells were scraped, collected and incubated on ice for 5 min. Cell lysates were centrifuged for 10 min at 2500 rpm at 4°C. The supernatants were aliquoted and stored at -80 °C until further processing. Lysate protein concentrations were measured using a bicinchoninic acid protein assay reagent kit (Thermo Scientific #23227). Protein samples were resolved on stain-free acrylamide gels (TGX Stain-Free FastCast Acrylamide kit, 10%, Bio-Rad, Mississauga, On) and transferred onto polyvinylidene fluoride (PVDF) membranes. Membranes were blocked with TBS-Tween 20 (0.1%) containing 3% bovine serum albumin (BSA) or dry milk, according to manufacturer instructions, for 1 h and incubated overnight at 4°C with the following primary antibodies: anti-E-cadherin (#3195; Cell Signaling Technology, Danvers, MA), anti-Cytokeratin 18 (#ab52948; Abcam, Cambridge, MA), anti-Cytokeratin 14 (#MS-115-P1ABX;

Thermo Scientific, Cheshire, UK), anti-α-Smooth muscle actin (#M0851; Dako, Glostrup, Denmark) and anti-p63 antibody (#sc-8431, Santa Cruz Biotechnologies, Dallas, TX). Bound primary antibody was detected using HRP-conjugated secondary antibodies (goat-anti-rabbit (#7074) or horse-anti-mouse (#7076); Cell Signaling Technology) followed by visualisation and quantification using a Bio-Rad ChemiDoc MP System (Bio-Rad Laboratories, Mississauga, On). Chemiluminescent signals were detected using Clarity western ECL substrate (Bio-Rad Laboratories) and analysed using Image Lab software (Bio-Rad Laboratories).

### 2.3.6.3 Three-dimensional embedded cells cultures

For 3D cultures, cells were embedded in solubilized basement membrane extract – Matrigel (Corning^®^ Growth Factor Reduced (GFR) Basement Membrane Matrix from Engelbreth-Holm-Swarm (EHS) mouse sarcoma) (CB40230C, Corning, NY, USA) at a cell density of 75 000 cells/100 μl of Matrigel. When required, Matrigel was diluted to different final concentrations by adding ice cold growth medium. Experiments were carried out in 35mm glass bottom poly-D lysine coated dishes, 14 mm microwell (P35GC-0-10-C, MatTEK Corporation, Ashland, MA, USA). Plates were manually evenly pre-coated on ice with 10 μl of Matrigel using the tip of a pipette. Dishes were left in an incubator at 37°C to allow the Matrigel to congeal for 30 minutes. For co-culture experiments, cell suspension from both cell types were counted and the proper number of each cell type mixed in the same tube. Cell suspension containing both cell types were centrifuged at 125G at room temperature for 7 minutes, the supernatant was removed and the tube was gently flicked to detach cells from the bottom of the centrifuge tube. The Matrigel was added directly to the cell pellet, gently mixed and the Matrigel-cells suspension was distributed rapidly in the microwells pre-coated with Matrigel. Dishes were incubated for another 30 minutes at 37 °C to allow Matrigel to congeal, then, the culture medium was added to cover the entire polymerized Matrigel/cell mix. The culture medium was changed every 2-3 days for 14 days.

### 2.3.6.4 Immunolabeling of 3D embedded cell cultures

Culture medium was aspirated and embedded cultures were rinsed twice with PBS. Cells were fixed in formaldehyde 4% for 10 minutes and permeabilized in PBS-Triton X-100 (0.5%) for 60 minutes. After being rinsed twice (10 minutes each) with PBS-Glycine (0.1%), cells were incubated in blocking solution (2% BSA dissolved in PBS with 0.1% of Tween 20 and 0.1% of Triton X-100). Primary antibodies were diluted in blocking solution and cells were incubated with primary antibody for 120 minutes at room temperature or overnight at 4°C (E-cadherin (#3195s) 1/200 (Cell Signaling, Beverly, MA, USA); α-Smooth-muscle-actin mouse mAb (M0851) 1/300 (Agilent, Santa Clara, CA, USA). Immunolabeling was followed by four washes with washing solution (PBS containing Tween20 (0.1%), BSA (0.1%) and Triton X-100 (0.5%)), for five minutes each. Cells were incubated with the appropriate secondary antibodies for 60 minutes (anti-rabbit IgG Alexa Fluor 488 (#4412s), anti-mouse IgG Alexa Fluor 555 (#4409s) both used at 1/1000 (Cell Signaling), or anti-rabbit IgG DyLight 488 (# 35552) used at 1/200 (ThermoFisher Scientific). Secondary antibody labeling was washed three times with washing solution and one time with PBS, for five minutes each. Nuclei were stained with 4', 6-diamidino-2-phenylindole (DAPI) in PBS. Embedded cultures were mounted with Fluoromount-G (0100-01, SouthernBiotech, Birmingham, AL, USA). Fluoromount-G was added in enough volume to cover the embedded cultures and fill up the microwell. Coverslip were added and mounted cultures were placed horizontally overnight at 4°C for 8 hours in the dark. Coverslips were sealed 24 hours after mounting with hot glue. Immunofluorescence images were obtained with a Nikon A1R+ confocal microscope (Nikon) and analyzed using NIS-elements software (Nikon).

### 2.3.6.5 Cryosections from embedded 3D cultures

Culture medium was removed and embedded cultures were washed twice with PBS, prewarmed at 37 °C. The matrix was detached from the microwell with a spatula, quickly placed on top of a layer of Tissue Freezing Medium (3801481, Leica Biosystems, Wetzlar, Germany) in a cryomold, and then covered with a second layer of Tissue Freezing Medium to fill the cryomold. The resulting block of cryomatrix containing the cells embedded in the matrix were placed at –80°C until use.

### 2.3.6.6 Masson's trichrome staining of cryosections

Embedded acini cryosections (8 μm) were fixed in Bouin’s solution overnight and then rinsed with water for 5 minutes. Sections were stained sequentially with Weigert’s iron hematoxylin for 10 minutes, Biebrich scarlet-acid fuchsin for 15 minutes, phosphomolybdic-phosphotungstic acid for 20 minutes and aniline blue (5 minutes), and washed with water between all coloration steps. Finally, the sections were treated with 1% acetic acid for 5 minutes, dehydrated in an alcohol series, cleared in xylene for 5 minutes and mounted using Permount (SP15100, Fisher Scientific, Burlington, ON, Canada). Slides were dried for a minimum of 4 hours after.

### 2.3.6.7 Immunolabeling of cryosections

Embedded acini cryosections (8 μm) were fixed in formaldehyde 4% for 15 minutes and blocked in 3% BSA dissolved in TBS-Tween20 (0.1%). Primary antibodies were diluted in TBS-Tween 0.1%, and sections were incubated with primary antibody for 60 minutes at room temperature. Sections were incubated with β-catenin (L54E2) mouse mAb (#2677s) 1/200 (Cell Signaling). Immunolabeling was followed by three washes with TBS-tween 0.1%. Sections were incubated with the secondary antibody (anti-mouse IgG Fab2 Alexa Fluor 647 (#4410s) used at 1/1000 (Cell Signaling). Nuclei were stained with DAPI in TBS-Tween 0.1%, and slides were mounted with Fluoromount-G (SouthernBiotech, 0100-01). Slides were placed at 4°C for 8 hours in the dark. Immunofluorescence images were obtained with a Nikon A1R+ confocal microscope (Nikon) and analyzed using NIS-elements software (Nikon).

## 2.3.7 Acknowledgments

We thank Calvin Roskelley (UBC) for the gift of SCp2 and SCg6 cells and Alicia Novel for her assistance. This work was supported by grant a Natural Sciences and Engineering Research Council grant (NSERC #418233-2012) to IP. IP is a recipient of a Fonds de Recherche du Québec-Santé (FRQS) and Quebec Breast Cancer Foundation career award and a Leader Founds from the Canadian Foundation for Innovation. AWO received a scholarship from La foundation Armand-Frappier. MB received scholarships from NSERC and from the Fonds de Recherche du Québec-Nature et technologie (FRQNT).

## 2.3.8 Author contribution

All experiments involving MCF-12A and Hs 578Bst cells were performed by AWO. MB performed the experiments with SCp2 and SCg6 cells. IP supervised the study and the progression of the project. AWO wrote the manuscript in collaboration with MB and IP.

## 2.3.10 Supplementary figures

**Supplementary figure 1.**
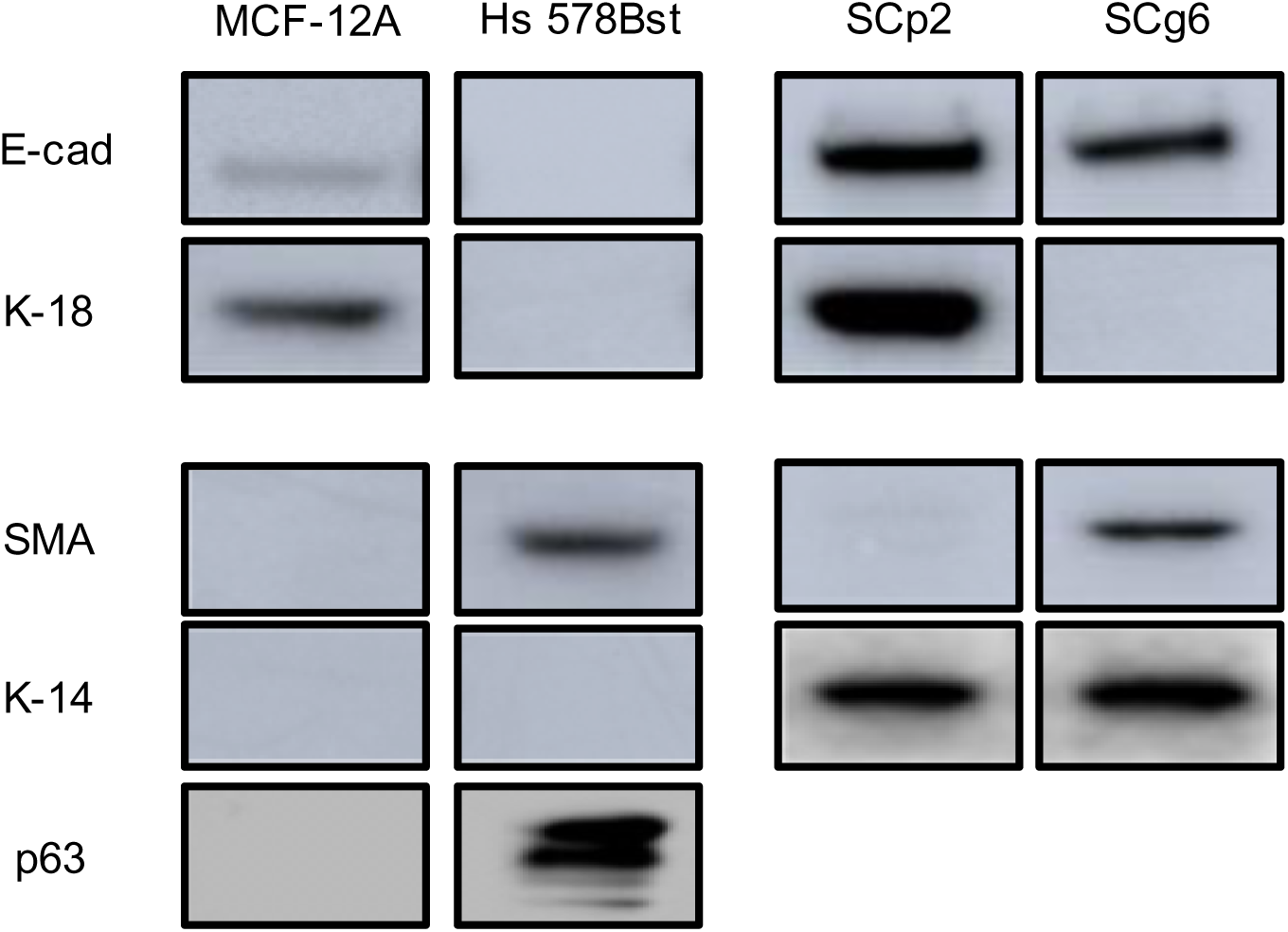
Human luminal epithelial MCF-12A and myoepithelial-like Hs 578Bst cells have more differentiated phenotypes than murine luminal epithelial SCp2 and myoepithelial SCg6 cells. MCF-12A, Hs 578Bst, SCp2 and SCg6 cells were cultured in 2D. Western blot analysis were performed using antibodies against E-cadherin (E-cad) and cytokeratin 18 (K-18), as well as SMA, Cytokeratin 14 (K14) and p63, markers of epithelial and myoepithelial cells, respectively. Results showed that in human cells, E-cad and K18 were present only in MCF-1CA, where as in mirine cells, E-cad was present in both SCp2 and SCg6. SMA and p63 were expressed only in myoepithelial cells, as expected, while K-14 was present in both SCp2 and SCg6.

**Supplementary figure 2.**
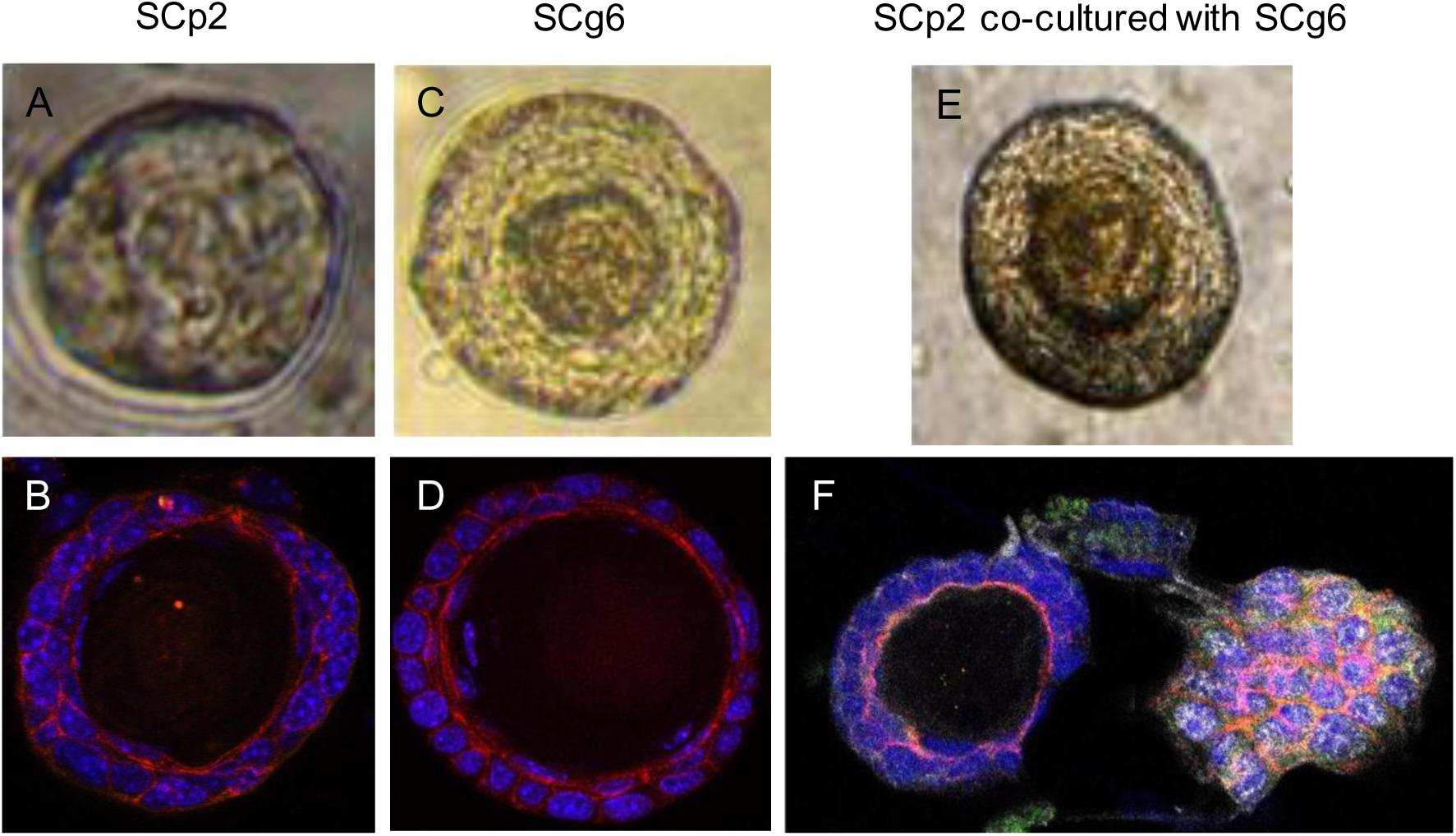
Co-cultures SCp2 and SCg6 cells did not form bilayered acini in Matrigel. (**A, C, E**) Representative single optical microscopy images of SCp2 cells (**A**), SCg6 cells (**B**) and SCp2 cells co-cultured with SCg6 cells (**E**) embedded in Matrigel. (**B, D, F**) Cells were immunolabeled with E-cadherin (red) and SMA (white). Nuclei are stained with DAPI (blue). SCg6 cells formed spheroids structures with lumen (**C, D**), similar to SCp2 (**A, B**). When SCp2 were co-cultured with SCg6 (**E, F**), some spheroids had a spheroids while other did not. SMA was present only on structures without lumen (**F**). SCp2 and SCg6 cells did not form bilayered spheroid with a lumen in co-culture.

## References

Anders M, Hansen R, Ding RX, Rauen KA, Bissell MJ, Korn WM (2003) Disruption of 3D tissue integrity facilitates adenovirus infection by deregulating the coxsackievirus and adenovirus receptor. Proc Natl Acad Sci U S A 100: 1943-8

Barsky SH, Karlin NJ (2005) Myoepithelial cells: autocrine and paracrine suppressors of breast cancer progression. J Mammary Gland Biol Neoplasia 10: 249-60

Birgersdotter A, Sandberg R, Ernberg I (2005) Gene expression perturbation in vitro‐‐a growing case for three-dimensional (3D) culture systems. Semin Cancer Biol 15: 405-12

Bissell MJ, Hall HG, Parry G (1982) How does the extracellular matrix direct gene expression? J Theor Biol 99: 31-68

Campbell JJ, Davidenko N, Caffarel MM, Cameron RE, Watson CJ (2011) A multifunctional 3D co-culture system for studies of mammary tissue morphogenesis and stem cell biology. PLoS One 6: e25661

Campbell JJ, Watson CJ (2009) Three-dimensional culture models of mammary gland. Organogenesis 5: 43-9

Carter EP, Gopsill JA, Gomm JJ, Jones JL, Grose RP (2017) A 3D in vitro model of the human breast duct: a method to unravel myoepithelial-luminal interactions in the progression of breast cancer. Breast Cancer Res 19: 50

Chitcholtan K, Asselin E, Parent S, Sykes PH, Evans JJ (2013) Differences in growth properties of endometrial cancer in three dimensional (3D) culture and 2D cell monolayer. Exp Cell Res 319: 75-87

Debnath J, Muthuswamy SK, Brugge JS (2003) Morphogenesis and oncogenesis of MCF-10A mammary epithelial acini grown in three-dimensional basement membrane cultures. Methods 30: 256-68

Desprez P, Roskelley C, Campisi J, Bissell M (1993) Isolation of functional cell lines from a mouse mammary epithelial cell strain: the importance of basement membrane and cell-cell interaction. Mol Cell Differ 1: 99-110

Deugnier MA, Moiseyeva EP, Thiery JP, Glukhova M (1995) Myoepithelial cell differentiation in the developing mammary gland: progressive acquisition of smooth muscle phenotype. Dev Dyn 204: 107-17

Dickson SR, Warburton MJ (1992) Enhanced synthesis of gelatinase and stromelysin by myoepithelial cells during involution of the rat mammary gland. J Histochem Cytochem 40: 697-703

Edmondson R, Broglie JJ, Adcock AF, Yang L (2014) Three-dimensional cell culture systems and their applications in drug discovery and cell-based biosensors. Assay Drug Dev Technol 12: 207-18

Ghajar CM, Bissell MJ (2008) Extracellular matrix control of mammary gland morphogenesis and tumorigenesis: insights from imaging. Histochem Cell Biol 130: 1105-18

Gudjonsson T, Adriance MC, Sternlicht MD, Petersen OW, Bissell MJ (2005) Myoepithelial cells: their origin and function in breast morphogenesis and neoplasia. J Mammary Gland Biol Neoplasia 10: 261-72

Gudjonsson T, Ronnov-Jessen L, Villadsen R, Rank F, Bissell MJ, Petersen OW (2002) Normal and tumor-derived myoepithelial cells differ in their ability to interact with luminal breast epithelial cells for polarity and basement membrane deposition. J Cell Sci 115: 39-50

Hassiotou F, Geddes D (2013) Anatomy of the human mammary gland: Current status of knowledge. Clin Anat 26: 29-48

Huh D, Hamilton GA, Ingber DE (2011) From 3D cell culture to organs-on-chips. Trends Cell Biol 21: 745-54

Ip MM, Darcy KM (1996) Three-dimensional mammary primary culture model systems. J Mammary Gland Biol Neoplasia 1: 91-110

Kaur G, Dufour JM (2012) Cell lines: Valuable tools or useless artifacts. Spermatogenesis 2: 1-5

Kenny PA, Lee GY, Myers CA, Neve RM, Semeiks JR, Spellman PT, Lorenz K, Lee EH, Barcellos-Hoff MH, Petersen OW, Gray JW, Bissell MJ (2007) The morphologies of breast cancer cell lines in three-dimensional assays correlate with their profiles of gene expression. Mol Oncol 1: 84-96

Krause S, Maffini MV, Soto AM, Sonnenschein C (2008) A novel 3D in vitro culture model to study stromal-epithelial interactions in the mammary gland. Tissue Eng Part C Methods 14: 261-71

Leblond CP, Inoue S (1989) Structure, composition, and assembly of basement membrane. Am J Anat 185: 367-90

Lee GY, Kenny PA, Lee EH, Bissell MJ (2007) Three-dimensional culture models of normal and malignant breast epithelial cells. Nat Methods 4: 359-65

Mabry KM, Payne SZ, Anseth KS (2016) Microarray analyses to quantify advantages of 2D and 3D hydrogel culture systems in maintaining the native valvular interstitial cell phenotype. Biomaterials 74: 31-41

Marchese S, Silva E (2012) Disruption of 3D MCF-12A breast cell cultures by estrogens‐‐an in vitro model for ER-mediated changes indicative of hormonal carcinogenesis. PLoS One 7: e45767

Pineda ET, Nerem RM, Ahsan T (2013) Differentiation patterns of embryonic stem cells in two-versus three-dimensional culture. Cells Tissues Organs 197: 399-410

Plante I, Laird DW (2008) Decreased levels of connexin43 result in impaired development of the mammary gland in a mouse model of oculodentodigital dysplasia. Dev Biol 318: 312-22

Radisky D, Hagios C, Bissell MJ (2001) Tumors are unique organs defined by abnormal signaling and context. Semin Cancer Biol 11: 87-95

Radisky ES, Radisky DC (2010) Matrix metalloproteinase-induced epithelial-mesenchymal transition in breast cancer. J Mammary Gland Biol Neoplasia 15: 201-12

Rudolph-Owen LA, Matrisian LM (1998) Matrix metalloproteinases in remodeling of the normal and neoplastic mammary gland. J Mammary Gland Biol Neoplasia 3: 177-89

Runswick SK, O'Hare MJ, Jones L, Streuli CH, Garrod DR (2001) Desmosomal adhesion regulates epithelial morphogenesis and cell positioning. Nat Cell Biol 3: 823-30

Sopel M (2010) The myoepithelial cell: its role in normal mammary glands and breast cancer. Folia Morphol (Warsz) 69: 1-14

Sternlicht MD, Kedeshian P, Shao ZM, Safarians S, Barsky SH (1997) The human myoepithelial cell is a natural tumor suppressor. Clin Cancer Res 3: 1949-58

Streuli CH, Schmidhauser C, Bailey N, Yurchenco P, Skubitz AP, Roskelley C, Bissell MJ (1995) Laminin mediates tissue-specific gene expression in mammary epithelia. J Cell Biol 129: 591-603

Su Y, Shankar K, Rahal O, Simmen RC (2011) Bidirectional signaling of mammary epithelium and stroma: implications for breast cancer‐‐preventive actions of dietary factors. J Nutr Biochem 22: 605-11

Talhouk RS, Mroue R, Mokalled M, Abi-Mosleh L, Nehme R, Ismail A, Khalil A, Zaatari M, El-Sabban ME (2008) Heterocellular interaction enhances recruitment of alpha and beta-catenins and ZO-2 into functional gap-junction complexes and induces gap junction-dependant differentiation of mammary epithelial cells. Exp Cell Res 314: 3275-91

Tibbitt MW, Anseth KS (2009) Hydrogels as extracellular matrix mimics for 3D cell culture. Biotechnol Bioeng 103: 655-63

Wang X, Sun L, Maffini MV, Soto A, Sonnenschein C, Kaplan DL (2010) A complex 3D human tissue culture system based on mammary stromal cells and silk scaffolds for modeling breast morphogenesis and function. Biomaterials 31: 3920-9

Warburton MJ, Mitchell D, Ormerod EJ, Rudland P (1982) Distribution of myoepithelial cells and basement membrane proteins in the resting, pregnant, lactating, and involuting rat mammary gland. J Histochem Cytochem 30: 667-76

Weigelt B, Bissell MJ (2008) Unraveling the microenvironmental influences on the normal mammary gland and breast cancer. Semin Cancer Biol 18: 311-21

Weigelt B, Ghajar CM, Bissell MJ (2014) The need for complex 3D culture models to unravel novel pathways and identify accurate biomarkers in breast cancer. Adv Drug Deliv Rev 69-70: 42-51

